# Species-specific developmental timing dictates expansion of the avian wing skeletal pattern

**DOI:** 10.1101/2021.03.22.436457

**Authors:** Holly Stainton, Matthew Towers

## Abstract

A fundamental question in biology is how species-specific rates of embryonic development are controlled in differently sized species. To address this problem, we compared wing development in the quail and the larger chick. We reveal that pattern formation (humerus to digits) is faster in the quail wing as determined by the earlier activation of 5’ *Hox* genes, termination of developmental organisers (*Shh* and *Fgf8*) and the laying down of the skeleton (*Sox9*). Using interspecies tissue grafts, we show that developmental timing can be reset during a critical window in which retinoic acid signalling is active. Accordingly, retinoic acid can switch developmental timing between the quail and chick, and also between the chick and the larger turkey. We show that the incremental growth rate is equivalent between all three species, suggesting that differences in the expansion of the skeletal pattern are governed primarily by the tempo of development. Our findings have implications for how development is timed both within and between different species.

## Introduction

Developmental timing can be defined as the rate at which embryos progress through a series of morphological states and sequential patterning events. Although we know much about how the embryo develops, our knowledge about how this sequence of events is timed between different species is currently not well understood ^(1, 2)^. Recent *in vitro* approaches have revealed that the rate of protein degradation in mouse cells is approximately twice as fast as is found in human cells, and that this correlates with the tempo of somitogenesis and motor neuron differentiation ^(3, 4)^. The avian wing provides an excellent *in vivo* system to understand species developmental timing, as we possess in-depth knowledge of the underlying mechanisms that pattern the proximo-distal axis (humerus to digits), and which rely on the integration of extrinsic signalling and autonomous timing processes ^(5)^. Thus, the specification of the chick wing skeletal pattern involves a switch from proximal signalling from the body wall (humerus/stylopod specification) to an autonomous timing mechanism operating in mesoderm cells at the distal tip of the outgrowing bud (digit/autopod specification) ^(6–6)^. Recent evidence suggests that this transition from proximal signalling to autonomous timing occurs during forearm/zeugopod specification ^(6, 10, 12)^. Retinoic acid emanating from the trunk of the embryo is implicated as the extrinsic signal involved in proximal specification ^(8, 9, 13–15)^. The specification of positional values that encode the different segments of the limb is associated with the progressive expression of genes encoding 5’ Hox proteins: *Hoxa/d10* provides a read-out of stylopod specification, and then *Hoxa/d11* followed by *Hoxa/d13* provide read-outs of zeugopod and autopod specification, respectively ^(16, 17)^.

Following proximo-distal specification, complex reciprocal epithelio-mesodermal signalling interactions sustain limb outgrowth as the population of *Sox9*-expressing condensing cartilage cells expand and the skeletal pattern is laid down. The undifferentiated distal mesoderm maintains the overlying apical ectodermal ridge by producing a signal encoded by the Bmp inhibitor, *Gremlin1* ^(18–18)^. The apical ectodermal ridge is a thickening of the distal epithelium that maintains limb outgrowth and is marked by the expression of *Fgf8* ^(21–21)^. However, the duration of proliferative growth of the chick wing is an autonomous property of the undifferentiated mesoderm and is controlled by the progressive Bmp-dependent decline in G1-S-phase entry, in both the distal tip and in the polarising region ^(7, 26)^ – a region of posterior-distal mesoderm that produces Sonic hedgehog (Shh), which is involved in specifying positional values along the antero-posterior axis (thumb to little finger) ^(27, 28)^. The laying down of the skeletal pattern along the proximo-distal axis is complete when *Sox9* is expressed in all condensing cartilage cells, proliferative outgrowth terminates in the distal mesoderm and the apical ectodermal ridge regresses ^(26)^.

Here we demonstrate that developmental timing is faster in smaller avian species and is associated with the tempo of *5’Hox* gene activation and the earlier laying down of the skeletal pattern. We implicate retinoic acid as the signal that sets the tempo of development, and we reveal that it is sufficient to switch quail to chick, and chick to turkey timing. We show that avian wings grow at a comparable rate, suggesting that expansion of the skeletal pattern is dictated by species differences in developmental timing.

## Results

### Proximo-distal patterning is faster in the quail compared to the chick wing

To understand how the tempo of development is controlled between differently sized species, we staged quail wings in reference to the Hamburger Hamilton (HH) staging system of the larger chick (Fig. 1a–c). The quail and chick are both in the Galliformes order and have incubation periods of 16 and 21 days, respectively. At day 3 of incubation, quail and chick embryos are at an equivalent stage (HH18/19), as determined by the appearance of the allantois (extra-embryonic membrane sac) and somites extending into the tail bud (approximately 36 pairs) ^(29–31)^. At this stage, the chick embryo is slightly wider than the quail embryo (between the wing buds), but it is not significantly different in length (from the tail bud to metencephalon – Supp Fig. 1). Assignment of the HH stage is based on the shape and gross morphology of the wing. Thus, at HH18/19, wing buds can be identified as slight symmetrical bulges protruding from the flank of the embryo (for the rest of the paper we define this as 0-hours - the last time-point that quail and chick wings are at an equivalent stage of development Fig. 1b,c). The full pattern of skeletal elements is laid down by HH29 when the apical ectodermal ridge regresses, which the chick wing reaches at 72 hours and quail wing reaches at 60 hours (Fig. 1b,c). Correspondingly, *Fgf8* is expressed in the chick wing apical ectodermal ridge for 72 hours and in the quail wing for 60 hours (Fig. 1d). In a similar manner, there is a 12-hour difference in the timing of *Shh* expression in quail and chick wing polarising regions. Thus, *Shh* expression persists for 36 hours in the quail and for 48 hours in the chick (until HH26 in both species, Fig. 1e). Furthermore, there is a 12-hour difference in the timing of *Sox9* expression, which is a marker of the expanding population of differentiating chondrogenic cells that prefigure the entire skeletal pattern by HH29 ^(32, 33)^. Thus, the onset of *Sox9* expression occurs at HH22 in both species, and this 12-hour difference in timing can be appreciated by its similar spatial pattern at HH27/28, which is reached at 48 hours in the quail and 60 hours in the chick (Fig. 1f). To further assess the timing of development, we analysed the anterior and posterior necrotic zones, which are regions of apoptosis in the wing ^(34, 35)^. As with the apical ectodermal ridge and differentiating chondrogenic cells, there is a 12-hour difference in the timing of apoptosis between quail and chick wings. Thus, the anterior necrotic zone persists between HH26 and HH27/28 and the posterior necrotic zone is detected at HH30 in both species (Fig. 1g).

**Figure 1.**
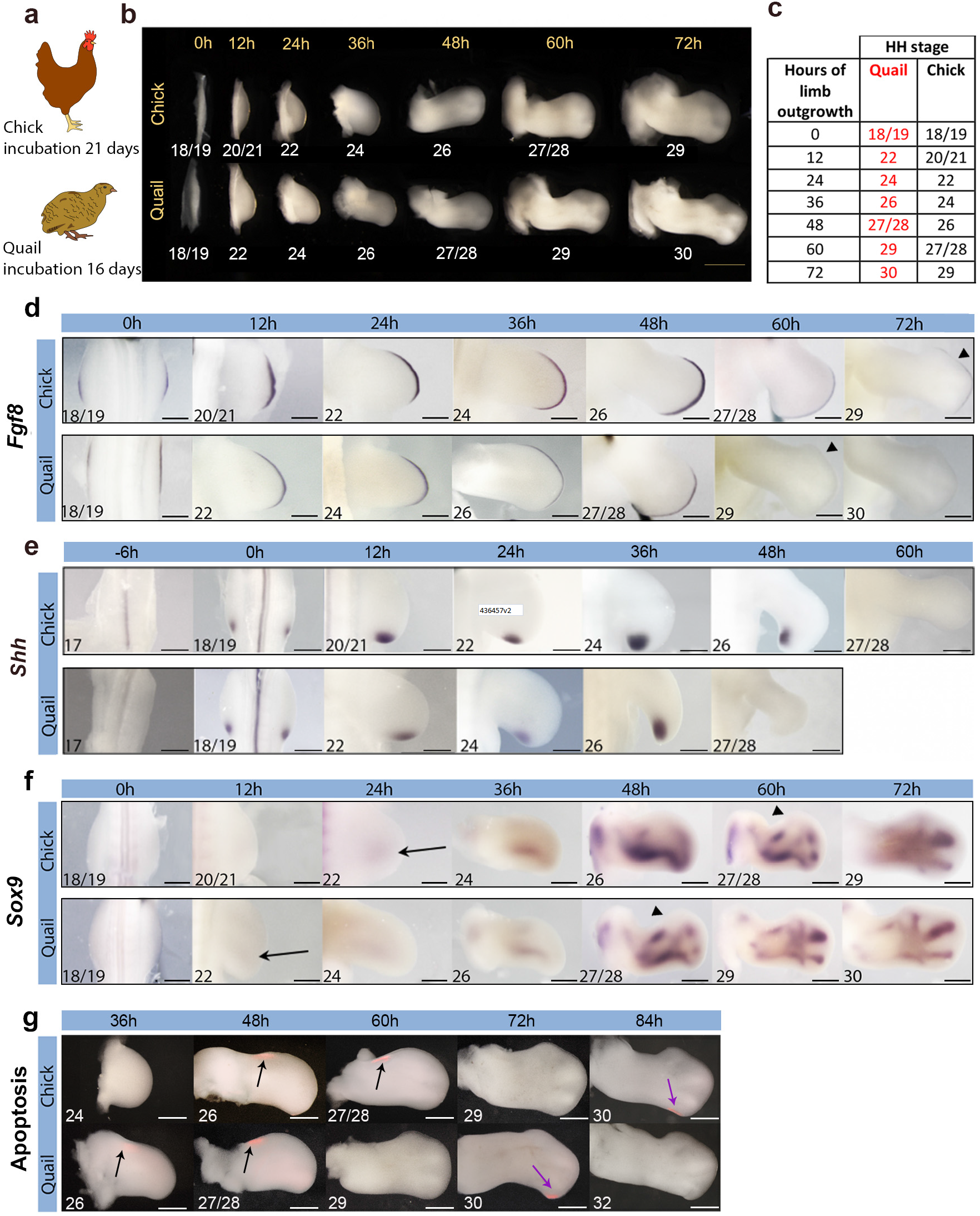
Developmental timing is accelerated in quail wing buds. **(a)** Schematics of the chick and quail that have 21- and 16-day incubation periods, respectively. **(b, c)** Hamburger Hamilton (HH) staging of chick and quail wings over 72 hours until HH29 and HH30, respectively - 0-hour refers to day 3 of incubation. (**d**) The apical ectodermal ridge marker, *Fgf8,* is down-regulated at HH29 - 72 hours in the chick wing and 60 hours in the quail wing, indicated by arrowheads - HH stage noted in the bottom left-hand corner. **(e)** *Shh* is expressed 12 hours longer in chick wings compared to quail wings until HH26 (HH stage noted in the bottom left corner). (**f**) Cartilage differentiation indicated by *Sox9* is 12 hours advanced in the quail wing compared to the chick (compare 48-hour quail wing and 60-hour chick wing arrowheads). Black arrows indicate the onset of *Sox9* expression at HH22. **(g)** Anterior (black arrows) and posterior (purple arrows) necrotic zones (red) are 12 hours advanced in quail wings compared to chick wings (*n* =4). Scale bars: a – 1mm, d – 12h, 24h −200μm; 36h −400μm; 0h, 48h, 60h −500μm; 72h −600μm. E - 12h, 24h - 300μm; 36h - 500μm; −6h, 0h, 48h, 60h - 600μm; 48h quail - 700μm. f – 12h, 24h −250μm; 0h −400μm; 36h −500μm; 48h, 60h −600μm; 72h −700μm. g– 36h, 48h, 60h −500μm; 72h −600μm, 84h −750μm.

These observations reveal that the developmental progression from HH18/19 to HH22 is faster in the quail wing bud, and this results in patterning being completed 12 hours earlier than it is in the chick wing (HH29 in both species).

### Proximo-distal specification is faster in quail wings compared to chick wings

Since the 12-hour difference in developmental timing between quail and chick wings is established between HH18/19 and HH22, we determined whether it is associated with the timing of proximo-distal positional value specification. The switch from proximal specification (stylopod) to intermediate specification (zeugopod) is indicated by the activation of *Hoxa11* expression, and the switch from intermediate specification (zeugopod) to distal specification (autopod) is indicated by the activation of *Hoxa13* expression ^(6, 12, 17, 36)^. Expression of *Hoxa11* is detectable in the quail wing bud at 6 hours and in the chick at 12 hours (Fig. 2a, HH20/21 in both cases), which resolves into a 12-hour difference in timing at later stages (compare 48-hour quail to 60-hour chick). By contrast, expression of *Hoxa13* is detectable in the quail wing bud at 12 hours and in the chick at 24 hours (Fig. 2b, HH22 in both cases), and this 12-hour difference in timing is maintained throughout outgrowth. Thus, the transition from stylopod to zeugopod specification occurs 6 hours later in the chick wing bud compared to the quail, and the transition from zeugopod to autopod specification occurs 12 hours later.

**Figure 2.**
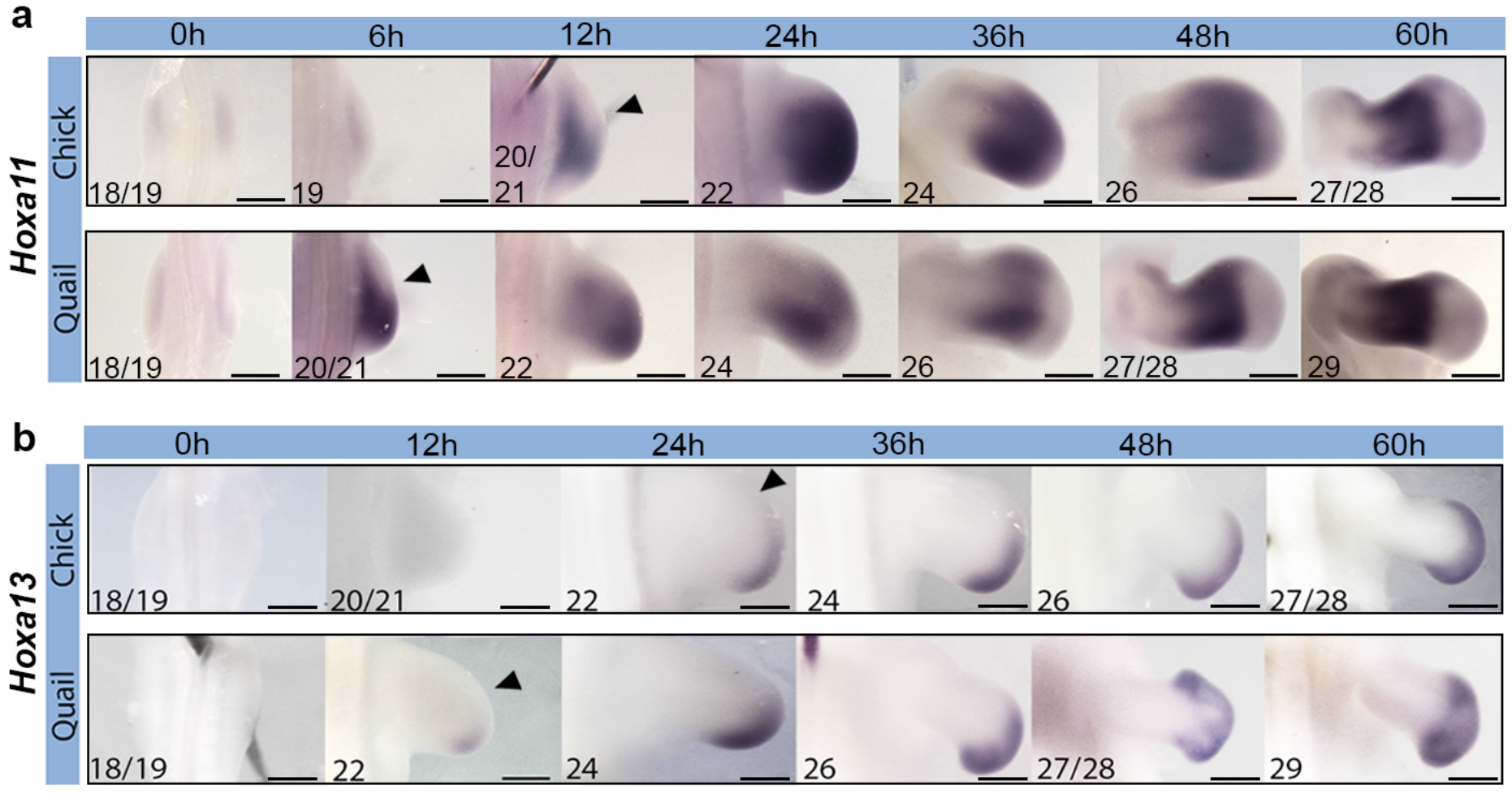
Proximo-distal specification is faster in quail wing buds. **(a)** *Hoxa11* is expressed at HH20/21, which is 6 hours earlier in quail compared to chick, arrowheads indicate onset of expression, HH stage noted in the bottom left corner. **(b)** *Hoxa13* is expressed at HH22, which is 12 hours earlier in quail compared to chick, arrowheads indicate onset of expression. Scale bars: a – 6h, 12h, 24h - 300μm; 0h, 36h - 500μm; 48h, 60h - 600μm; b – 12h, 24h - 200μm; 0h, 48h chick - 300μm; 36h - 400μm, 48h quail - 600μm, 60h - 800μm.

These findings reveal that proximo-distal positional value specification is 12 hours faster in the quail wing bud compared to the chick.

### Stability and resetting of developmental timing in interspecies grafts

To gain insights into how the 12-hour difference in developmental timing is set between quail and chick wings between HH18/19 and HH22, we performed a series of reciprocal tissue grafting experiments to ascertain if species developmental timing is reset or maintained. We chose the polarising region, as it expresses *Shh* for a species-specific duration, and regulates its cell cycle parameters for an autonomously timed duration in the chick wing (Fig. 1e) ^(11, 37)^.

We performed interspecies polarising region grafts to the anterior margins of host wing buds at 12 hours when developmental timing is advanced in the quail compared to chick (HH21 chick and HH22 quail – Fig. 3a,c, note that stage-matched intraspecies control grafts maintain their normal duration of *Shh* expression – Supp Fig. 2). We found that grafts performed at these stages maintain their species timing of *Shh* expression. Thus, at 48 hours, *Shh* expression is undetectable in quail cells grafted to a chick wing but is detectable in the host (Fig. 3b). In the reciprocal experiment, *Shh* expression is detectable in chick cells grafted to a quail wing but is undetectable in the host (Fig. 3d). In addition, both grafted quail and chick wing bud polarising region cells maintain species-specific cell cycle parameters typical of their donor age (Fig. 3e; 63.3% G1-phase cells in quail graft vs. 71% in chick host; 72.6 chick graft vs. 64% in quail host). Rather than being an autonomous process, it has also been suggested that the termination of *Shh* expression requires the displacement of *Gremlin1-*expressing cells by a critical distance from the polarising region in order to break down a self-propagating extrinsic signalling loop ^(38)^. However, in chick wings that received quail polarising region grafts, *Shh* expression is terminated in donor cells independently of their proximity to the duplicate domain of *Gremlin1* expression in the host (Supp Fig. 3). Thus, these findings show that the timing of both *Shh* expression and cell cycle parameters are autonomously determined after HH21.

**Figure 3.**
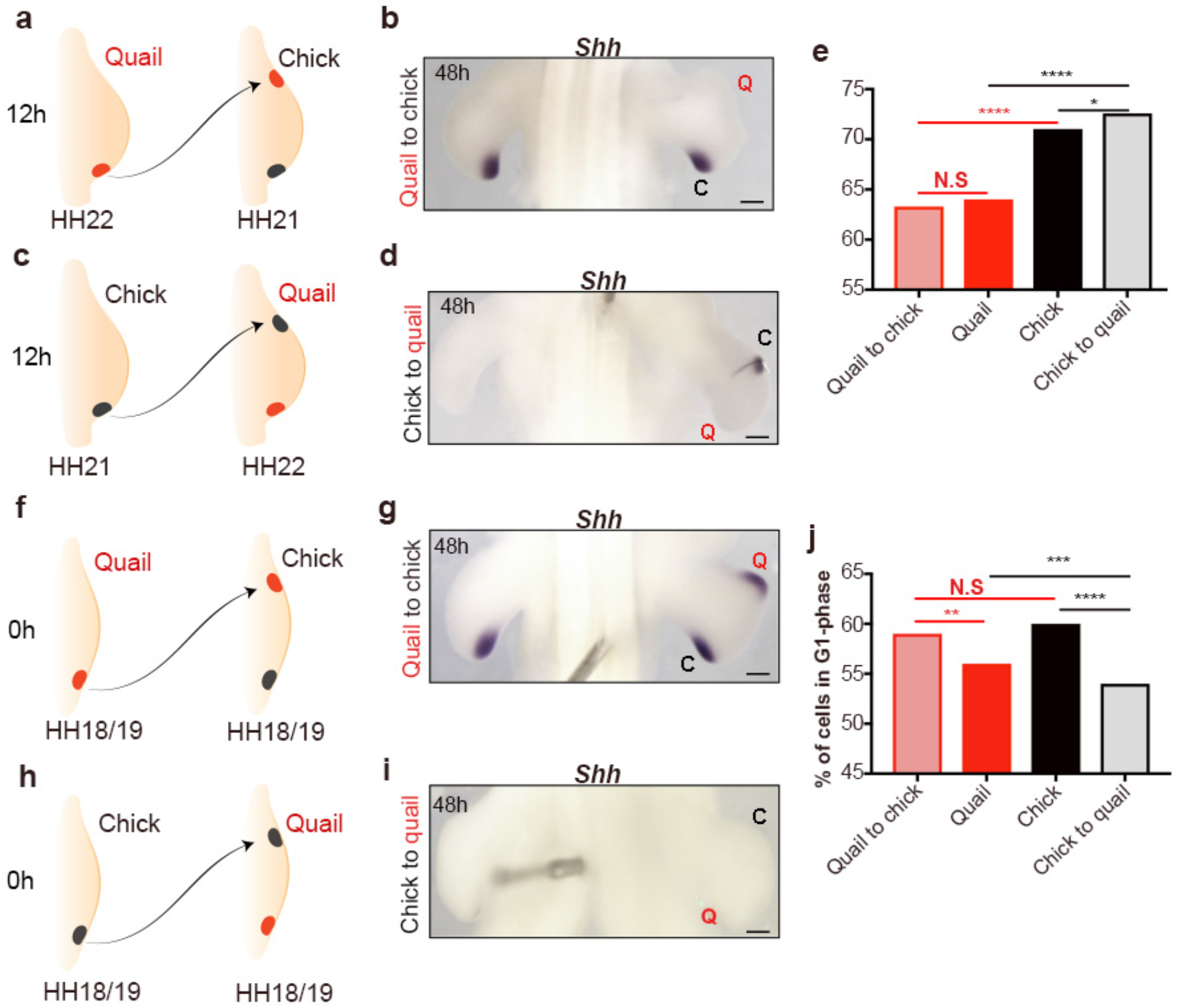
Resetting potential of species developmental timing in polarising region grafts. In interspecies polarising region grafts between 12-hour quail (HH22) and chick (HH21) wings (made to the anterior margins of host wing buds) **(a,c),** *Shh* expression is maintained according to donor timing **(b** *n*=7/10, **d** *n*=5/7). **(e)** Cell cycle parameters in grafts are close to donor values but are significantly different from the host polarising region 24 hours after the graft was performed (Pearson’s *χ*2 test) *p*-values = quail vs. quail grafted to chick = 0.24, chick vs. quail grafted to chick = 0.00001, chick vs. chick grafted to quail = 0.013, quail vs. chick grafted to quail = 0.00001. In interspecies polarising region grafts between 0-hour quail and chick wings (HH18/19 in both species) **(f, h)**, *Shh* expression is reset according to host timing **(g**, *n*=9/10, **i** n=3/5). **(j)** Cell cycle parameters in grafts are reset close to host values 24 hours after the graft was performed. (Pearson’s *χ*2 test) *p-*values = quail vs. quail grafted to chick = 0.02, chick vs. quail grafted to chick = 0.09, chick vs. chick grafted to quail = 0.00001, quail vs. chick grafted to quail = 0.00012. *p-*values *–* N.S = >0.05 ***= <0.001 ****= <0.0001. Scale bars: - 500μm

We also performed interspecies polarising region grafts at 0 hours when developmental timing is equivalent in chick and quail wing buds (HH18/19 - Fig. 3f,h). Unlike with the 12-hour grafts, *Shh* expression is reset to the host timing in 0-hour interspecies grafts. Thus, at the 48-hour time-point, *Shh* expression duration is prolonged in quail cells in a chick wing, and is prematurely terminated in chick cells in a quail wing (Fig. 3g,i, respectively - note that 0-hour intraspecies control grafts maintain their normal duration of *Shh* expression - Supp Fig. 3). In addition, cell cycle parameters are also reset close to host values in both grafted quail and chick wing bud polarising regions (Fig. 3j; 59% G1-phase cells in quail graft vs. 60% in chick host; 54% in chick graft vs. 56% in quail host).

These results reveal that *Shh* expression and cell cycle parameters are autonomously maintained in 12-hour polarising region grafts when quail and chick developmental timing is different (HH21 vs. HH22), but that they can be reset in 0-hour grafts when developmental timing is equivalent (HH18/19).

### Retinoic acid can reset the species-specific timing of *Shh* expression

Since the timing of *Shh* expression can be reset by the host environment in interspecies polarising region grafts performed at 0 hours, but not at 12 hours, this suggests the existence of a transient resetting signal. Retinoic acid is the only known signal in the chick wing bud that, although present at high-levels at 0 hours (HH18/19), becomes cleared from the distal region of the wing bud by 12 hours (HH21) (14, 39-41), as indicated by down-regulation of its downstream transcriptional target *Meis1,* and the up-regulation of the gene encoding the retinoic acid degrading enzyme, Cyp26b1^(10, 42)^ (Fig. 4). The loss of distal *Meis1* expression occurs earlier in the quail wing bud at approximately 6 hours, compared to 12 hours in the chick wing bud (Fig. 4a – HH20/21 in both species). However, as with 5’ *Hox* gene expression, the spatial expression of *Meis1* resolves into a 12-hour difference by HH22, which is reached at 12 and 24 hours in quail and chick wing buds, respectively (Fig. 4a). Additionally, between 0-12 hours of wing bud outgrowth, *Cyp26b1* levels rise significantly faster in the quail compared to the chick (Fig. 4b). Therefore, this finding indicates an increased rate of retinoic acid degradation in the wing buds of the smaller species.

**Figure 4.**
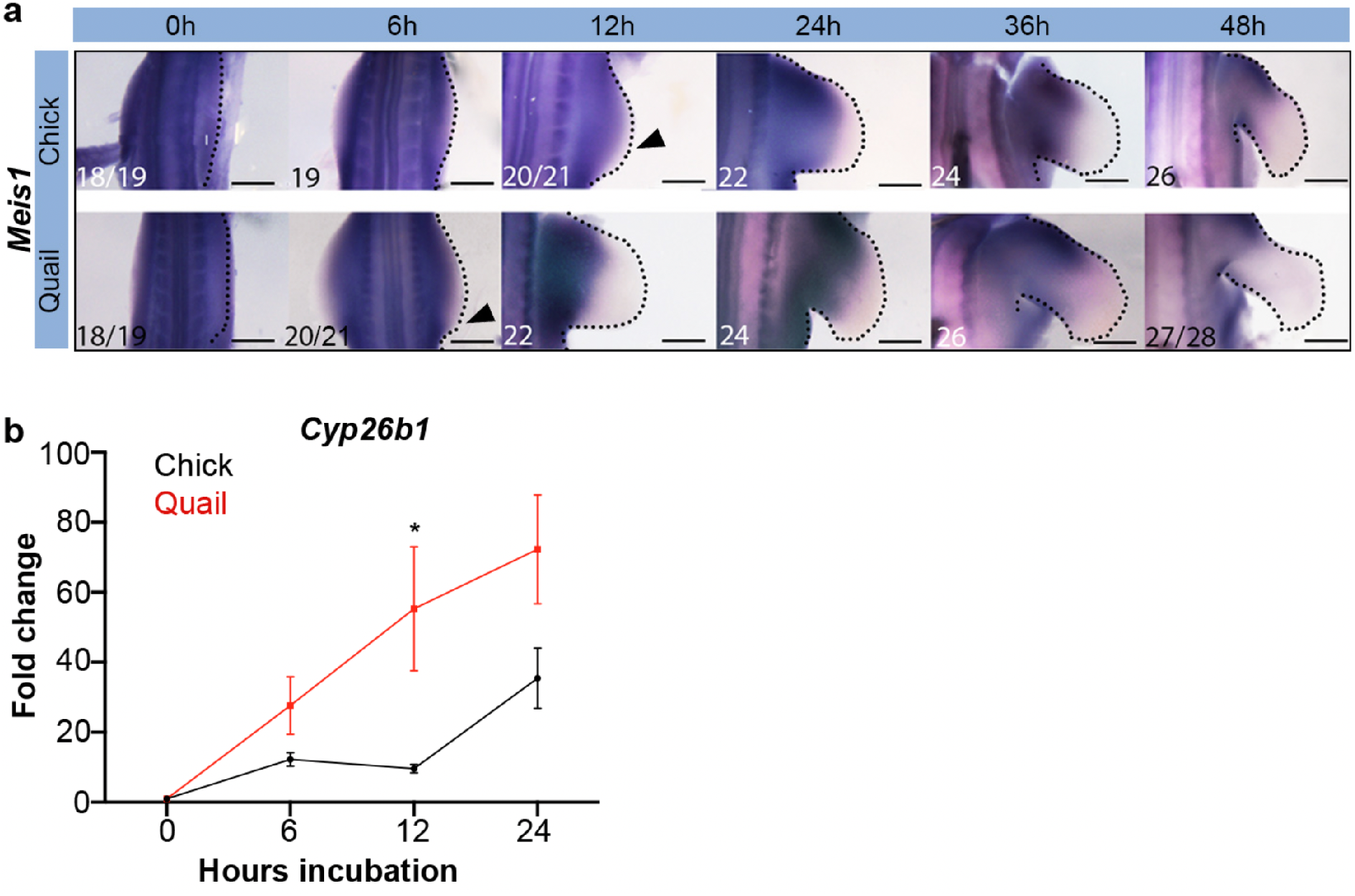
Retinoic acid degradation occurs earlier in quail wing buds. **a**) *Meis1* expression is down-regulated in the distal part of the wing at 12 hours in the chick and 6 hours in the quail, as indicated by arrowheads. **b**) A qPCR time-course reveals a significantly higher fold-change in *Cyp26b1* expression in 12-hour quail wing buds compared to chick wing buds (*p*-value=0.028) - the fold changes at 6 and 24 hours are not significantly different between species (*p-*value = 0.08 and 0.11, respectively). Student unpaired *t*-tests were performed on *n*=3 (quail) and *n*=4 (chick) repeats of 10 pooled wing buds. *p-*values: *=<0.05 Scale bars: a - 12h, 24h - 300μm; 0h, 6h - 400μm; 36h - 500μm; 48h - 650μm

The differences in the expression of genes that regulate and respond to retinoic acid signalling between quail and chick wing buds could suggest that it is a signal that can set developmental timing. Therefore, we asked if the transient maintenance of retinoic acid signalling by carrier beads for 12-20 hours in the chick wing bud ^(43, 44)^ would reverse the autonomy of *Shh* expression timing in quail polarising region grafts made to the chick wing at the 12-hour time-point (Fig. 5a). Indeed, by transiently prolonging retinoic acid signalling, *Shh* expression timing is reset in the quail polarising region graft, as it is maintained for approximately the same duration as is seen in the host chick polarising region (approx. 56 hours - Fig. 5b, compare with failure to reset timing in the same experiment minus retinoic acid - Fig, 3b, d). It is worth noting that the duration of *Shh* expression is also extended in the retinoic acid-treated host chick polarising region (compare to level of residual *Shh* expression in the contralateral untreated wing in Fig. 5b). These observations show that retinoic acid can reset the species-specific timing of *Shh* expression.

**Figure 5.**
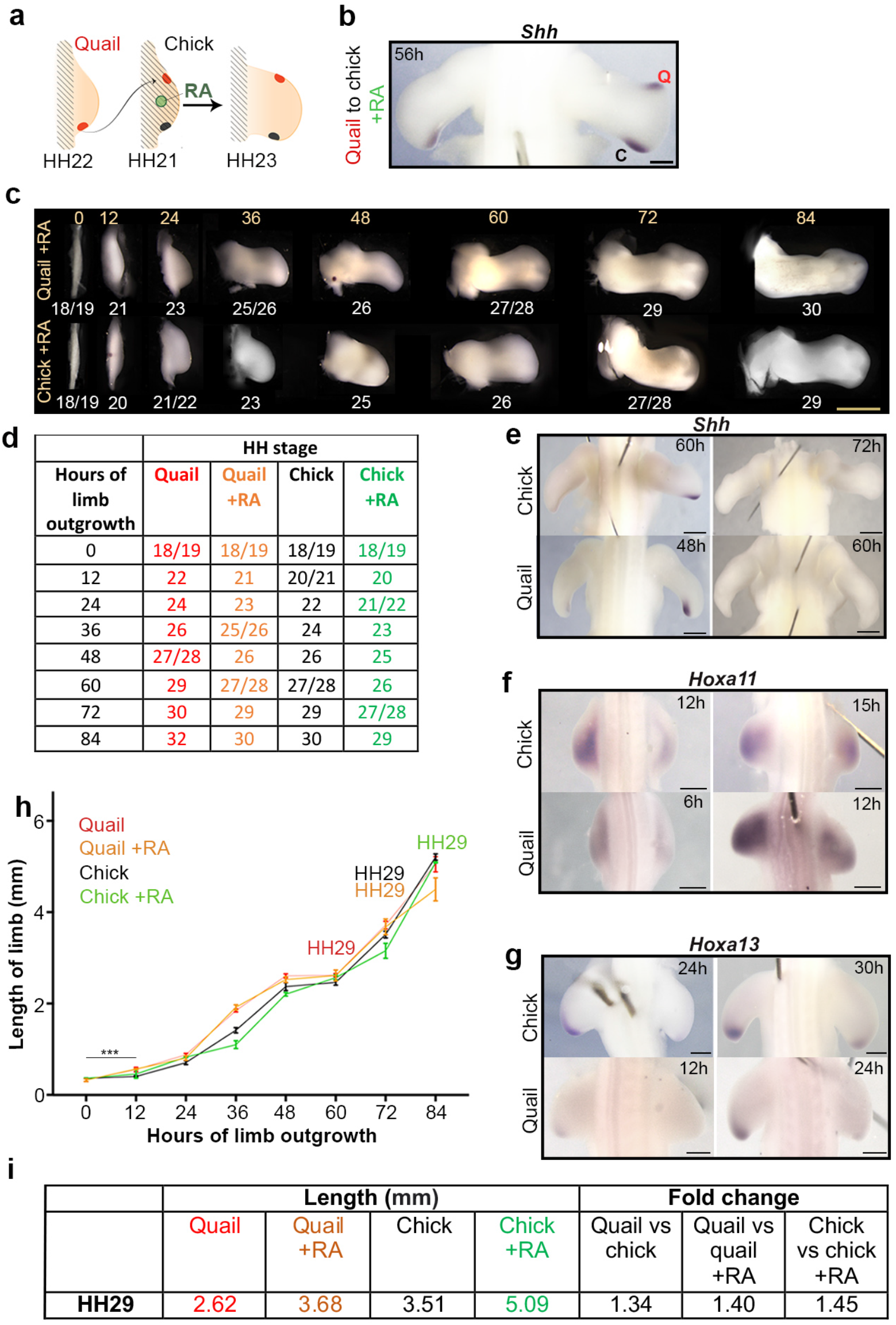
Retinoic acid can set species developmental timing. **(a)** HH19 chick wings were treated with retinoic acid-soaked beads (green circle), and at 12 hours (HH21) received polarising regions grafts from 12-hour quails (HH22); note, black hatching indicates retinoic acid. **(b)** *Shh* expression is prolonged until approximately 56 hours in both the host and donor (*n*=3/4). **(c, d)** HH stages of chick and quail wing buds treated with retinoic acid at 0 hours, compared with the contralateral untreated wing. **(e)** Retinoic acid-treated wings express *Shh* until 60 hours in the chick (*n*= 4/4) and 48 hours in the quail (*n*=3/3), compared to 48 and 36 hours in control untreated. **(f)** Retinoic acid-treated wings express *Hoxa11* at 15 hours in the chick (*n*=3/5), and 12 hours in the quail (*n*=3/5), compared to 12 and 6 hours in control untreated. **(g)** Retinoic acid-treated wings express *Hoxa13* at 36 hours in the chick (*n*=3/4) and 24 hours in the quail (*n*=3/5), compared to 24 and 12 hours in control untreated wings. **(h)** Quail wings grow at a significantly faster rate compared to chick wings between 0-12 hours, as determined by student *t*-test (*p*-value = 0.0008) (*n*=7). However, there is no significant difference in incremental changes in growth rates from 12-72 hours as determined by Wilcoxon tests (*p*-value = 0.688 - *n*=6-14). After 12 hours, Wilcoxon tests also reveal no significant difference in incremental growth rates between chick vs. chick +RA, chick vs. quail +RA, chick +RA vs. quail, quail vs. quail +RA, and chick +RA vs. quail +RA (*p* - values = >0.99, >0.99, 0.563, 0.438, 0.688 respectively). *n*= 4-16. **(i)** Lengths of quail, retinoic acid-treated quail, chick and retinoic acid-treated chick wings and fold differences at HH29. *p-*values: *=<0.05, **=<0.01 Scale bars: b - 500μm; d, - 1mm; e, f - 400μm, g, h –250μm,

### Retinoic acid can set species developmental timing

Having demonstrated that retinoic acid can reset the 12-hour difference in the timing of *Shh* expression between the quail and chick, we questioned if it could have a general role in setting development timing in the wing. Indeed, when retinoic acid is applied on beads to HH18/19 quail and chick wing buds at 0 hours to transiently maintain it for 12-20 hours ^(43, 44)^, the subsequent developmental progression through HH stages occurs approximately 12 hours later than normal, such that HH29 is reached at 72 hours and 84 hours, respectively (Fig. 5c,d). Correspondingly, as predicted from the polarising region grafting experiments (Fig. 5a,b), retinoic acid extends the duration of *Shh* expression for 12 hours in both quail and chick wings (until 48 hours and 60 hours, respectively Fig. 5e), which is HH26 in both species. In a similar manner, prolonged retinoic acid exposure also delays zeugopod specification (*Hoxa11*) for 3 hours in chick wing buds and for 6 hours in quail wing buds (Fig. 5f – HH20/21), and delays autopod specification *(Hoxa13*) for 12 hours in both species (Fig. 5g – HH22). However, consistent with recent studies, inhibiting retinoic acid does not precociously activate *Hoxa13,* due to autopod specification also being controlled by an autonomous epigenetic programme ^(10, 37)^ (Supp Fig. 4). These results show that prolonged retinoic acid treatment slows the development of both quail and chick wings. An implication of this finding is that retinoic acid-treated quail wings develop comparably to chick wings.

### Quail and chick wings have equivalent incremental growth rates

Having shown that developmental timing is faster in quail wings compared to chick wings, we asked if this is associated with changes in the rate of elongation along the proximo-distal axis. Therefore, we measured quail and chick wings from the trunk to the distal tip every 12 hours. We also measured the lengths of retinoic acid-treated quail and chick wings. Analyses of the data reveal that after 12 hours the incremental growth rates are not significantly different between species, and also between retinoic acid-treated and untreated wing buds (Fig. 5h). Therefore, the rate of growth is equivalent between quail and chick wings and is not linked with the tempo of development. A significant implication of this finding is that the length of the wing at HH29 varies considerably between quail and chick wings and those treated with retinoic acid. Therefore, quail wings treated with retinoic acid are 1.4-fold longer than untreated quail wings, and similarly, chick wings are 1.34-fold longer than quail wings (Fig. 5i).

### Chick wings treated with retinoic acid have turkey developmental timing

Having revealed that retinoic acid-treated quail wings and untreated chick wings have equivalent developmental timing (Fig. 5), we asked if retinoic acid-treated chick wings replicate timing found in a larger species. Therefore, we chose the turkey, which like the quail and chick, belongs to the Galliformes order (Fig. 6a –note the turkey has an incubation period of 28 days).

**Figure 6.**
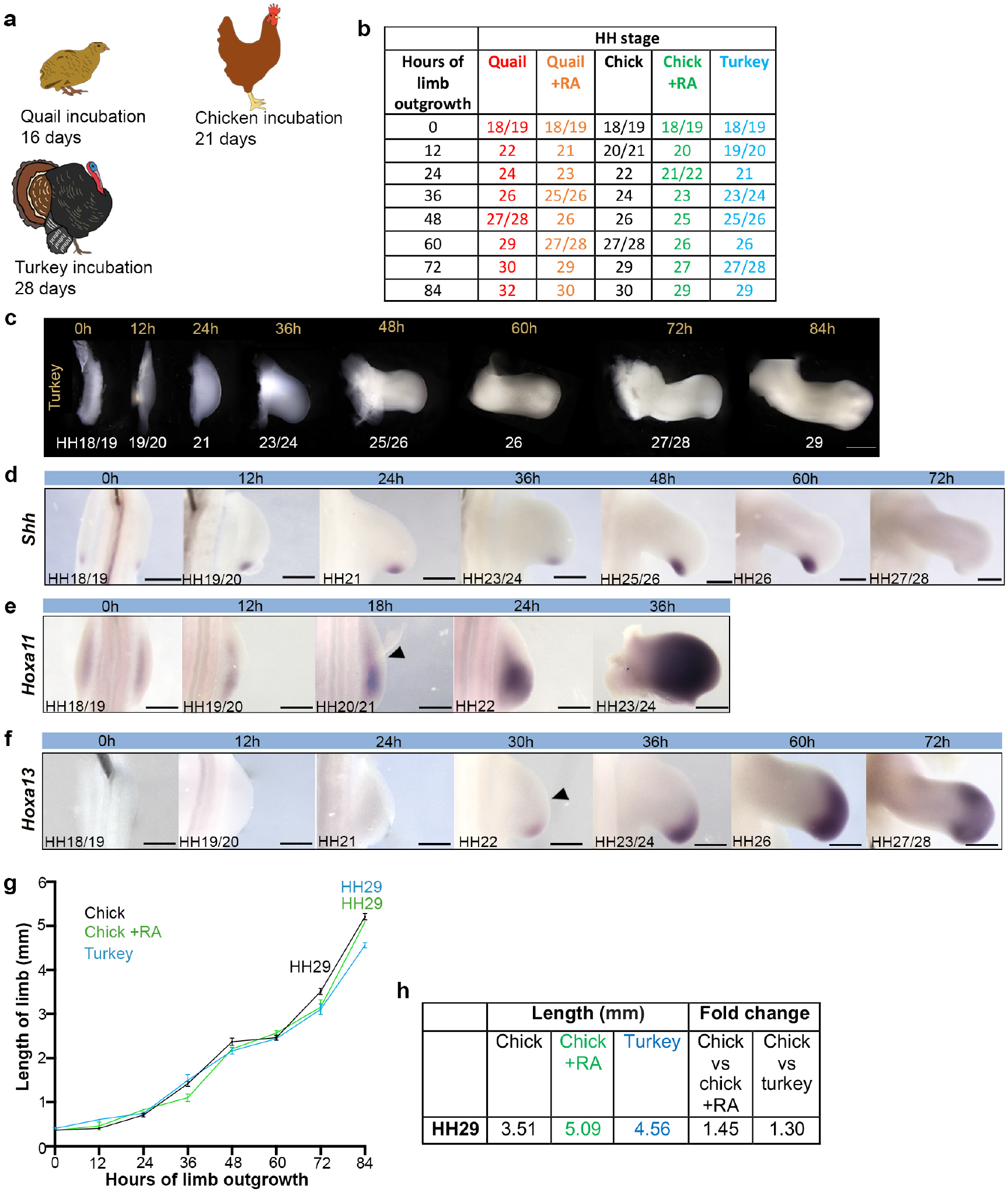
Retinoic acid treated chick wings replicate turkey developmental timing and pattern scaling. **(a)** Schematics of quail, chick and turkey that have 16, 21 and 28-day incubation periods, respectively. **(b,c)** HH staging of turkey wings over 84 hours until HH29 - 0-hour refers to day 4 of incubation. **(d)** *Shh* is downregulated at HH26, HH stages noted in the bottom left of each panel. **(e)** *Hoxa11* is expressed at HH20/21, arrowheads indicate onset of expression, and **(f)** *Hoxa13* is expressed at HH22, arrowheads indicate onset of expression. **(g)** Proximo-distal lengths of chick, chick +RA and turkey wings buds until 84 hours (HH29 in turkey and chick +RA wings). Wilcoxon tests reveal no significant difference in incremental growth rates between turkey vs. chick +RA, and turkey vs. chick (*p*-values = >0.437, 0.219, respectively). *n*= 4-14. Note, Wilcoxon tests also reveal no significant difference in incremental growth rates between turkey vs. quail +RA, turkey vs. quail, *p*-values = >0.99, 0.563, respectively). **(h)** Lengths of chick, retinoic acid-treated chick (Chick +RA), and turkey wings and fold differences at HH29. Scale bars: c - 750μm; d - 500μm, e-0-36h - 500μm, 60h, 72h - 600μm f-0h, 24h - 400μm, 12h, 36h-72h-500μm

We staged turkey wings according to the HH staging system of the chick, starting at HH18/19, which is reached at day 4 of incubation (note that the quail and chick reach HH18/19 at day 3). At HH18/19 the turkey embryo is significantly longer than the quail, but not the chick (from the tail bud to metencephalon), and is similar in width to both quail and chick embryos (between the wing buds, Supp Fig. 5a,b). During the next 12 hours, turkey wing buds progress to HH19/20, whereas chick wing buds reach HH20/21, and subsequently, the developmental timing of HH stage progression resolves into a 12-hour difference between the two species by 48 hours. Thus, HH29 is reached in 84 hours in turkey wings compared to 72 hours and 60 hours for chick and quail wings, respectively (Fig. 6b,c).

To investigate developmental timing further in the turkey wing, we analysed the expression of *Shh,* and found that transcripts can be detected for 60 hours (until HH26) in turkey wings (Fig. 6d). This is comparable to the duration of *Shh* expression in retinoic acid-treated chick wing buds (Fig. 5e), which is 12 hours longer than is found in untreated chick wings (48 hours – Fig. 1e). We also determined the timing of proximo-distal specification in turkey wing buds, by analysing 5’ *Hox* expression. The onset of *Hoxa11* expression (zeugopod specification) occurs at 18 hours (HH20/21), and *Hoxa13* expression (autopod specification) at 30 hours (HH22) (Fig. 6e,f, respectively), which are comparable timings to those found in retinoic acid-treated chick wing buds (Fig. 5g,h).

We also measured the incremental rate of growth along the proximo-distal axis and found that it is equivalent between turkey wings, untreated chick wings and retinoic acid-treated chick wings (Fig. 6g). Thus, at HH29, turkey wings are 1.3-fold longer than chick wings and, similarly, retinoic acid-treated chick wings are 1.45-fold longer than chick wings (Fig. 6h). These results show that retinoic acid-treated chick wings and turkey wings have equivalent developmental timing and rates of growth.

## Discussion

We have described a new mechanism that explains how the pace of embryonic wing development is controlled between different avian species (Fig. 7). The duration of stylopod and zeugopod specification (red and green) is variable (12 to 30 hours). Coinciding with the onset of autopod specification (blue), the autonomously timed programme of distal development (white) then continues for a similar duration until patterning is complete when the skeletal elements have been laid down. However, because the rate of growth is equivalent, species differences in developmental timing influence expansion and scaling of the skeletal pattern (Fig. 7). Our interspecies grafting experiments implicated retinoic acid as the signal that sets the tempo of development. Indeed, transiently prolonging retinoic acid signalling slows down the tempo of 5’*Hox* gene activation and therefore quail wings develop comparably to chick wings (Fig. 7), and chick wings develop comparably to turkey wings (Fig. 7).

**Figure 7.**
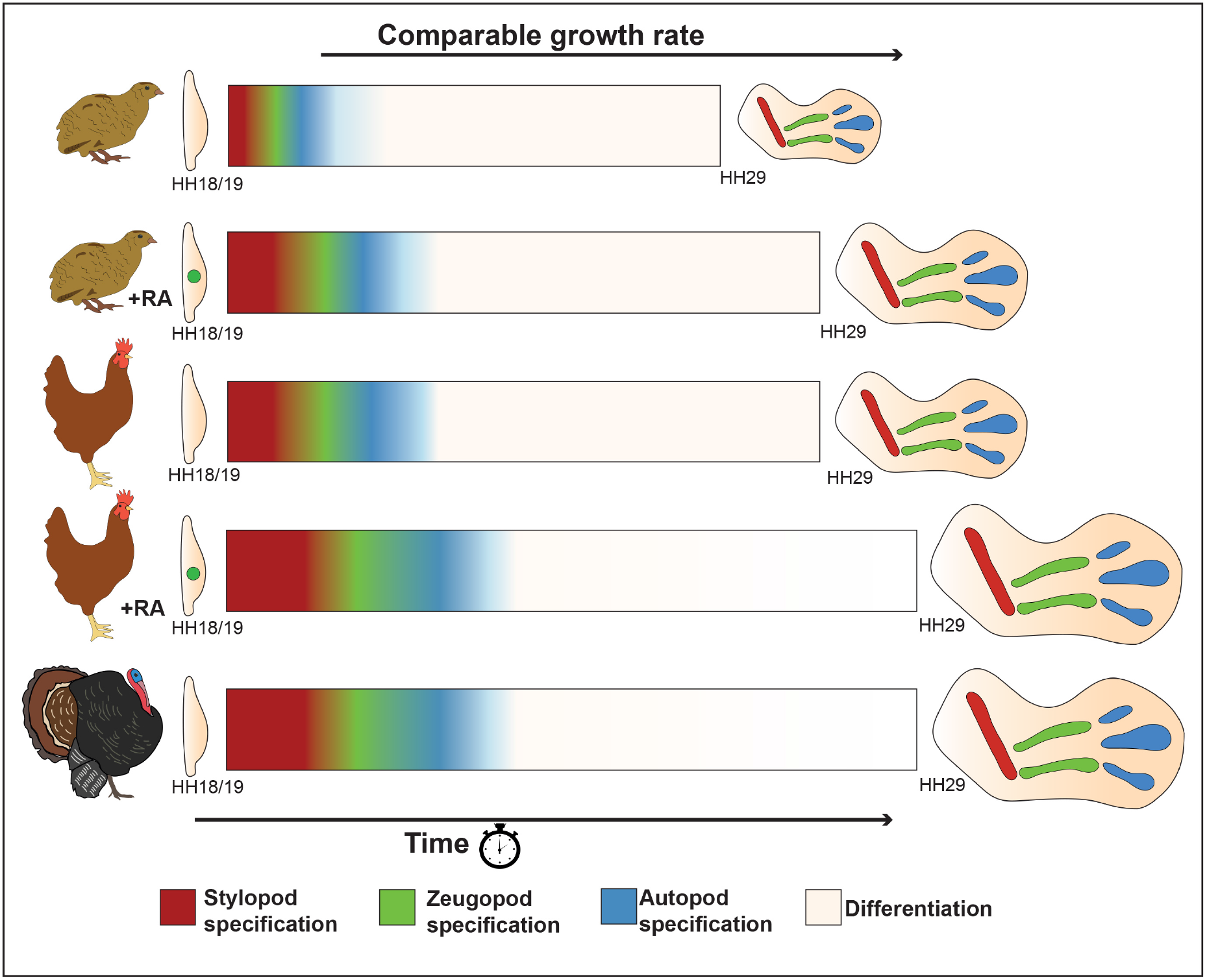
Developmental timing scales avian wing dictates expansion of the avian wing skeletal pattern. Schematics depicting the timing of proximo-distal specification and differentiation in avian wings (quail, chick and turkey) from HH18/19 until the end of the patterning phase at HH29, when the skeletal elements have been laid down. In this model, the stylopod (red) is specified by high levels of retinoic acid signalling; intermediate specification of the zeugopod (green) by low levels retinoic acid signalling at HH20/21 and autonomous timing, and distal specification of the autopod (blue) triggered by an autonomously timed epigenetic mechanism and the loss of retinoic acid signalling by HH22. The duration of stylopod and zeugopod specification (red and green) varies between species, however the duration of the autonomous programme (blue and white) remains relatively constant. Thus, the timing of development alongside a comparable growth rate results in species differences in the expansion of skeletal progenitor cells. Consequently, at HH29 when the complete skeletal pattern is laid down there is a 1.34-fold difference in the size of quail and chick wings, a 1.3-fold difference in the size of chick and turkey wings, and a 1.75-fold difference in the size of quail and turkey wings (schematics of HH29 wings are scaled appropriately).

We provided insights into the underlying mechanism that determines the variable species duration of proximo-distal specification (Fig. 7). The degradation of a proximal signal considered to be retinoic acid - is interpreted into a gradient of Meis1, which creates a permissive environment for the activation of 5’ *Hox* expression ^(12)^. Opposing Fgf signals from the apical ectodermal ridge also influence the distribution of retinoic acid from the flank of the embryo ^(12, 13, 42)^. In this model, high Meis1 levels are associated with stylopod specification (Hoxa10, red – Fig. 7); low Meis1, zeugopod specification (Hoxa11, green - Fig. 7), and absent Meis1, autopod specification (Hoxa13, blue – Fig. 7). Quail, chick and turkey embryos have similar trunk widths; therefore, our data indicate that this parameter does not influence the distribution of retinoic acid in the wing bud. Instead, we implicated the retinoic acid-degrading enzyme, Cyb26b1, in timing the removal of retinoic acid in the wing bud and setting the tempo of 5’ *Hox* gene activation. Thus, *Cyp26b1* expression in the quail wing bud increases at a significantly faster rate compared to the chick. In addition, the earlier depletion of retinoic acid signalling in the quail wing bud is indicated by the precocious loss of the distal domain of *Meis1* expression. Therefore, species-specific rates of retinoic acid degradation are associated with the tempo of 5’*Hox* gene expression and proximo-distal specification.

Following the variable period of stylopod and zeugopod specification in avian wings (red, green), the autonomously timed programme of distal development continues for a similar duration until patterning is complete (white - Fig. 7). The autonomous programme is triggered by the depletion of retinoic acid^(6, 10, 12)^ and co-ordinates the developmental timing of autopod specification (blue, Fig. 7), proliferation, differentiation, apoptosis, and organiser duration (white, Fig. 7 - note *Hoxa13* is expressed until at least day 10 preventing the duration of autopod specification from being be determined). Since the autonomous programme (autopod specification) is activated later in larger species, and is not linked to incremental rate of growth, this influences the size of the avian wing skeleton (Fig. 7) - a 1.75-fold difference in length between the quail and turkey wing at HH29.

We previously demonstrated that the duration of the autonomous programme is caused by the Bmp-dependent decline in proliferation rates in the distal mesoderm and that this is associated with the duration of both the polarising region ^(11, 37)^ and the apical ectodermal ridge ^(26)^. Therefore, it is likely that the cell cycle constitutes an overarching developmental timing mechanism, because it is intimately coupled with the other autonomously regulated processes – differentiation and apoptosis. Indeed, evidence that the cell cycle could constitute a developmental timer arises from the similarities between the chick wing and cultured oligodendroctye progenitor cells. In both cases, retinoic acid is implicated in triggering the onset of a cell cycle timer that involves the progressive lengthening of the G1-phase of the cell cycle (37, 45-47), and is associated with the activation of D-cyclin-dependent kinase inhibitors, which are important negative regulators of the G1-S phase transition ^(11, 47)^. Having shown that the autonomous programme runs for the approximately same duration in avian wings, it will be interesting to determine its temporal parameters in the limbs of non-avian species.

The widespread distribution of retinoic acid in the embryo could suggest that it has a general role in developmental timing. For instance, retinoic acid promotes the expression of anterior *Hoxb* genes along the main body axis and is removed by Cyp26b1 to permit the expression of posterior *Hoxb* genes^(48)^. Thus, the relative timing of *Hoxb* expression is suggested to underlie evolutionary changes in avian limb position ^(48, 49)^. However, it remains to be determined if retinoic acid affects developmental timing independently of growth along the main body axis, as we suggest it does in the limb. Nevertheless, these considerations could suggest that retinoic acid coordinates developmental timing throughout the embryo.

## Methods

### Avian husbandry and wing measurements

Bovans Brown chicken eggs (*Gallus gallus domesticus*), Japanese quail eggs (*Coturnix japonica*), and Bronze turkey eggs (*Meleagris gallopavo domesticus*) were incubated at 37°C and staged according to the Hamburger Hamilton staging system ^(30)^ based on the number of somites present, and by characteristic morphological features of the wing bud. HH18/19 is reached by incubation day 3 in quails and chicks, and day 4 in turkeys, and is referred to in this study as 0 hours of wing outgrowth. Embryos of the appropriate age were dissected in PBS and measurements of the proximo-distal axis were taken down the centre of the limb bud, from the proximal boundary of the limb with the body wall, to the distal tip of the limb bud, accounting for elbow bend where appropriate.

### Whole mount *in situ* hybridisation

Embryos were fixed in 4% PFA overnight at 4°C then dehydrated in methanol overnight at −20°C. Embryos were then rehydrated through a methanol/PBS series, washed in PBS, then treated with proteinase K for 20 mins (10 μg/ml−1), washed in PBS, fixed for 30 mins in 4% PFA at room temperature and then prehybridised at 67°C for 2 hours (50% formamide/50% 2x SSC). 1 μg of antisense DIG-labelled mRNA probes were added to 1 ml of hybridisation buffer (50% formamide/50% 2× SSC) at 67°C overnight. Embryos were washed twice in hybridisation buffer, twice in 50:50 hybridisation buffer and MAB buffer, and then twice in MAB buffer, before being transferred to blocking buffer (2% blocking reagent 20% foetal bovine serum in MAB buffer) for 3 hours at room temperature. Embryos were transferred to blocking buffer containing anti-digoxigenin antibody (1:2000) at 4°C overnight, then washed in MAB buffer overnight before being transferred to NTM buffer containing NBT/BCIP and mRNA distribution visualised using a LeicaMZ16F microscope.

### Flow cytometry

Polarising regions or distal mesenchyme pooled from 8-12 replicate experiments were dissected in PBS under a LeicaMZ16F microscope using fine surgical scissors, and digested into single cell suspensions with trypsin (0.05%, Gibco) for 30 mins at room temperature. Cells were briefly washed in PBS, fixed in 70% ethanol overnight, washed in PBS and re-suspended in PBS containing 0.1% Triton X-100, 50 μg/ml^−1^ of propidium iodide and 50 μg/ml^−1^ of RNase A (Sigma). Dissociated cells were left at room temperature for 20 mins, cell aggregates were removed by filtration and single cells analysed for DNA content with a FACSCalibur flow cytometer and FlowJo software (Tree star Inc.). Based on ploidy values cells were assigned G1, S, or G2/M phases, and this was expressed as a percentage of the total cell number (5,000–12,000 cells in each case). Statistical significance of numbers of cells in different phases of the cell cycle (G1 vs. S, G2 and M) between pools of dissected wing bud polarising region tissue (12–15 in each pool) was determined by Pearson’s χ^2^ tests to obtain two-tailed *P*-values (significantly different being a *p*-value of less than 0.05 – as in ^(37)^).

### Apoptosis analysis

Whole chick and quail wing buds were dissected in PBS and transferred to Lysotracker (Life Technologies, L-7528) PBS solution (1:1000) in the dark pre-warmed to 37°C. Wing buds were incubated for 1 hour at 37°C, washed in PBS, and fixed overnight in 4% PFA at 4°C. Wing buds were then washed in PBS and progressively dehydrated through a methanol series.

### Polarising region grafts

Polarising region grafts were performed as described in ^(50)^. Briefly, donor embryos were dissected in PBS and the polarising regions removed using sharpened tungsten needles then transferred to the host embryo where they were grafted to equivalently sized regions of the host anterior limb bud and held in place with 25μm diameter platinum pins.

### Quantitative PCR (qPCR)

Ten whole limb buds at 0, 6, 12 and 24 hours were dissected from either quail or chick embryos. Total RNA was extracted using TRIzol Reagent (Life Technologies), purified using a Direct-zol RNA kit (Zymo Research) and cDNA prepared using SuperScript III Reverse Transcriptase (Invitrogen). qPCR was performed on an Applied Biosystems StepOne RTPCR machine using SYBR Green Master Mix (Thermo Fisher Scientific) and a primer set for *Cyp26b1* was designed against a sequence which was present in both chicken and quails, spanning exon junctions (Thermo Fisher Scientific). 5 ng cDNA was used per reaction (20μl volume) with cycle conditions of 95 °C for 20 sec, followed by 32 cycles of 95 °C for 1 sec and 60 °C for 20 sec. All reactions were carried out in triplicate and average C_T_ values normalized against Eukaryotic *18S rRNA* Endogenous Control expression (Thermo Fisher Scientific). Unpaired student *t*‐tests compared the mean relative expression, and measured significance of expression change between appropriate samples. Applied Biosystems StepOne Software V2.3 was used to analyse the data and GraphPad Prism8 used to construct graphs.

### Bead implantation

Sieved AGX1-2 beads (150 or 200 μm in diameter, Sigma) were soaked in a stable form of all-*trans*-retinoic acid, TTNPB (Sigma, 0.05 mg/ml dissolved in DMSO, Sigma) or AGN193109 (Sigma, 1 mg/ml dissolved in DMSO, Sigma) for 1 hour and then washed in DMEM before being grafted to the middle of wing buds using a sharp tungsten needle. TTNPB has been shown to diffuse from AGX1-2 beads over an approximate 12-20-hour period and can be used to model RA distribution in chick wing buds due to comparable patterning effects, kinetics and diffusion constants ^(40, 43, 44)^.

### Data availability statement

The authors declare that the data supporting the findings of this study are available within the paper and its supplementary information files. Original data files including gating strategy related to the flow cytometry data in figure 3 can be found in Data S1 (flow cytometry source data) in the supplementary information file.

## Acknowledgements

We acknowledge Cheryll Tickle and Marysia Placzek for critical reading

## Author contributions

HS performed and analysed the experiments and co-wrote the paper with MT who devised the study

**The authors declare no competing interests**

