## supplementary data for "Species-specific developmental timing dictates expansion of the avian wing skeletal pattern"

Figs. S1 to S5

Data S1 (Flow cytometry source data)

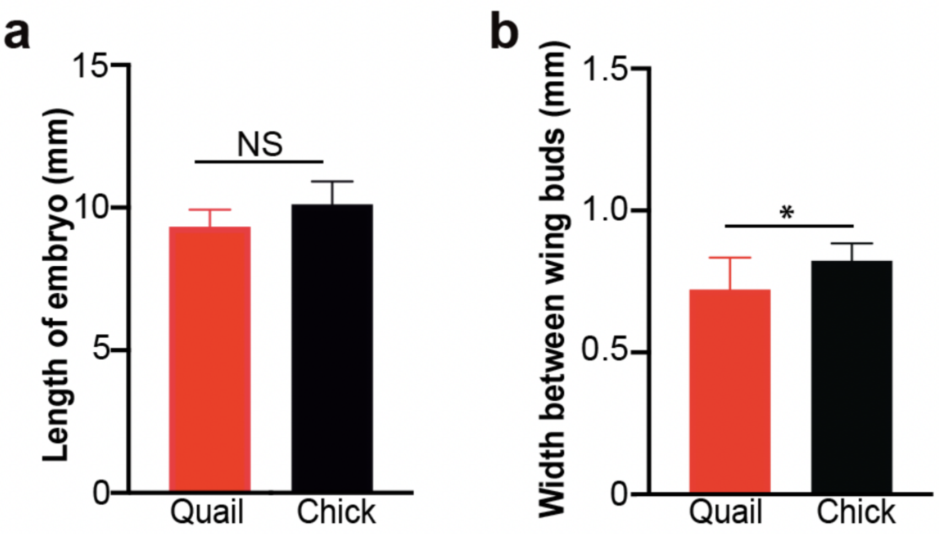


**Supplementary Figure 1. Lengths and widths of quail and chick embryos at HH18/19**

**(a)** Lengths of quail and chick embryos at 0 hours (HH18/19 – tail bud to metencephalon) are not significantly different, as indicated by student *t*-tests (*p*-value =0.055 *n*=6 and 10). **(b)** Widths of quail and chick embryos (between wing buds) at 0 hours are statistically different as indicated by student *t*-tests (*p*-value = 0.033, *n*=6 and 10). *p*-values: *=<0.05

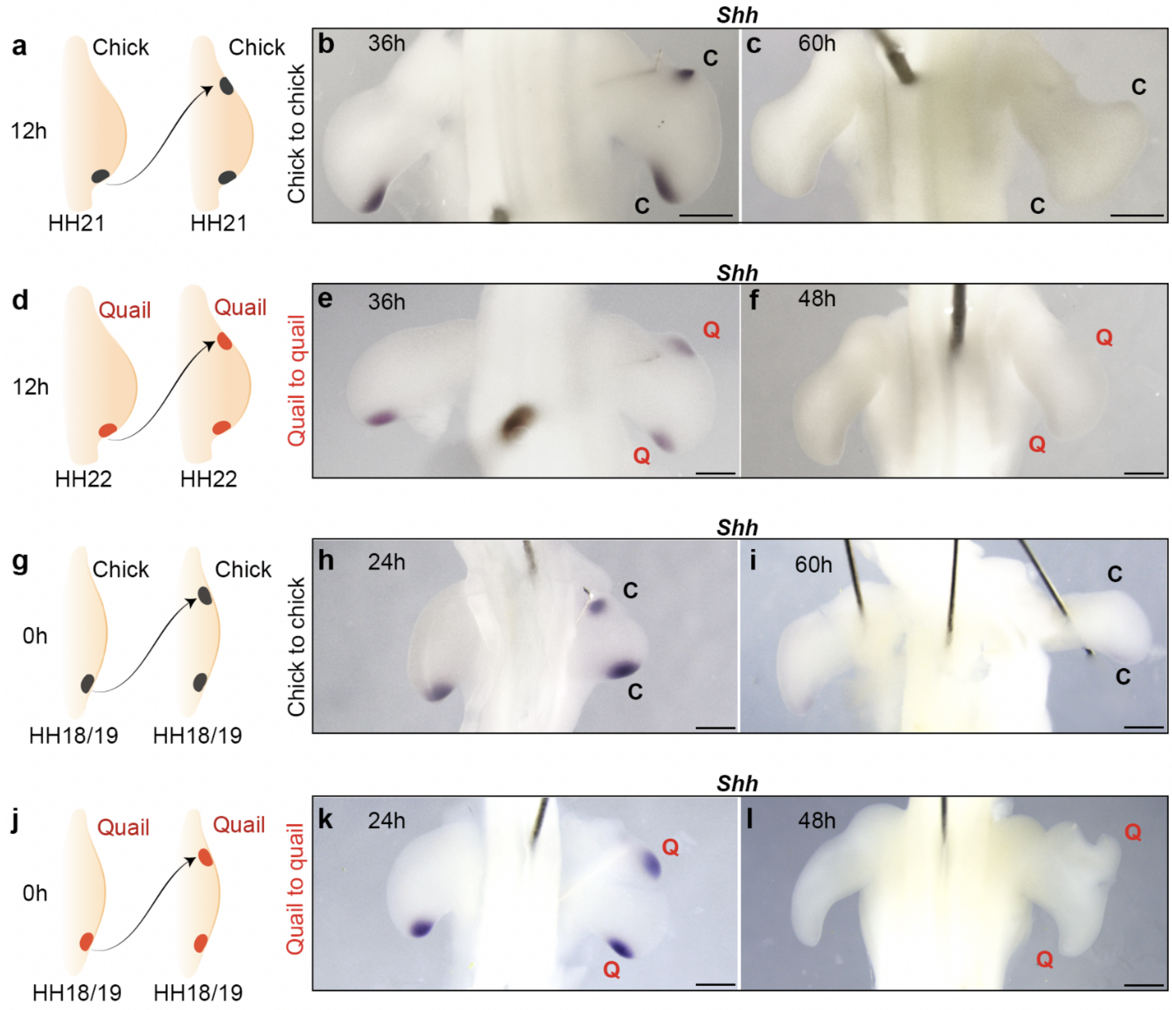


**Supplementary Figure 2. *Shh* maintains its duration in intraspecies polarising region grafts**

Control intraspecies chick (HH21) **(a-c)** and quail (HH22) **(d-f)** polarising region grafts made to the anterior margins of host wing buds at 12h. *Shh* is expressed at 36 h **(b,** *n*=4/5**, e** *n*=6/6**)** and terminates at the correct time as shown at 60h **(c** *n*=4/4), and 48h **(f** *n*=5/6). Control intraspecies chick **(g-i)** and quail **(j-l)**, HH18/19, polarising region grafts made to the anterior margins of host wing buds at 0h. *Shh* is expressed at 36 h **(h,** *n*=3/3**, k** *n*=4/4**)** and terminates at the correct time as shown at 60h **(i** *n*=3/4), and 48h **(l** *n*=3/3).

Scale bars**:** h,k = 300μm; f,b,e = 500μm c,i,l = 700μm

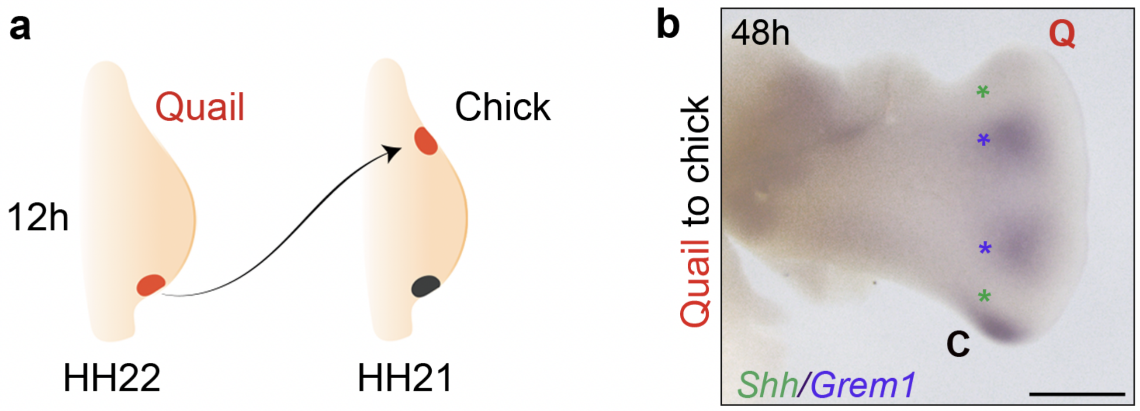


**Supplementary Figure 3. Termination of *Shh* is intrinsically controlled**

**(a)** Polarising regions grafted from 12-hour quail wing buds (HH22) to the anterior margins of chick wing buds (HH21). **(b)** Expression of *Shh* is terminated in the quail cells (upper green asterisk) and is observed in endogenous chick polarising region cells (lower green asterisk) at 48 hours. Both chick and grafted quail wing polarising regions induce a domain of *Grem1* expression in adjacent chick cells (purple asterisks): *Grem1* expression (lower purple asterisk) is adjacent to endogenous chick *Shh* expression, and a duplicated domain of *Grem1* expression is observed (upper purple asterisk) adjacent to where quail *Shh* would have been expressed. The loss of quail *Shh* expression demonstrates that *Grem1* expressing cells do not need to be displaced a critical distance by growth in order for *Shh* expression to be terminated at the correct time [(1)]. Note equivalent distance between *Grem1* domains and anterior (403.3µm) and posterior (398.3µm) margins, *p*-value = 0.7944 indicated by Student *t*-tests. (*n*= 3/3).

Scale bar = 600µm

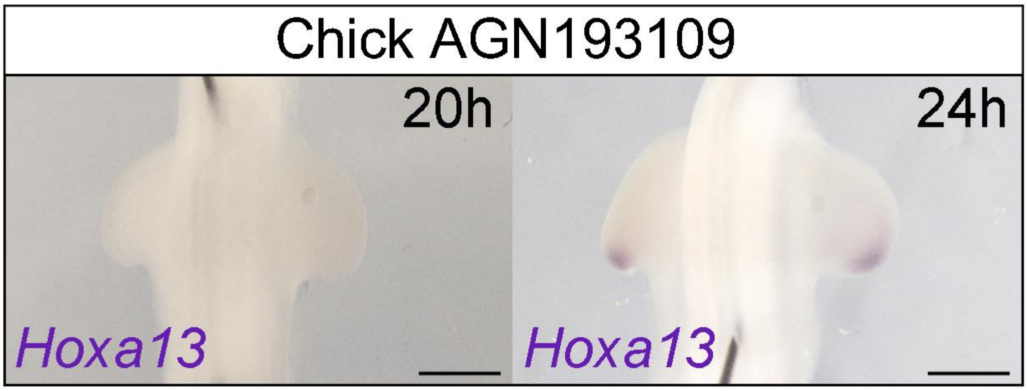


**Supplementary Figure 4. Retinoic acid inhibition does not precociously activate *Hoxa13* expression**

Right-hand chick wing buds were treated with AGN193109 at HH19 and compared to control untreated left wings. The onset of *Hoxa13* expression occurs normally at 24h hours (HH22) in both treated and untreated wings (*n*=6/6). Scale bar = 500µm

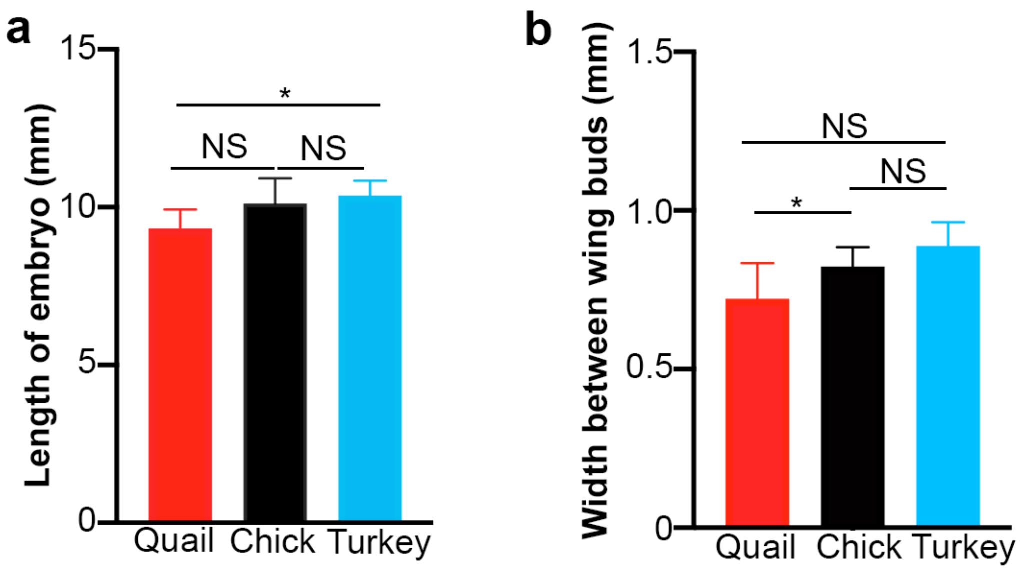


**Supplementary Figure 5 . Retinoic acid-treated chick wings develop according to turkey timing**

**(a)** Lengths of turkey and chick embryos at 0 hours (HH18/19 – tail bud to metencephalon) are not significantly different (*p*-value =0.63), however turkeys are significantly longer than quail embryos (*p*-value=0.036), as indicated by student *t*-tests. **(b)** Width of turkey embryos (between wing buds) at 0 hours are not significantly different to chick embryos (*p*-value=0.140), or quail embryos (*p*-value=0.056), as indicated by student *t*-tests (*n*=3, 6 and 10 – turkey, quail, chick, respectively). *p*-values: *=<0.05

### 0hr polarising region grafts

#### Quail control

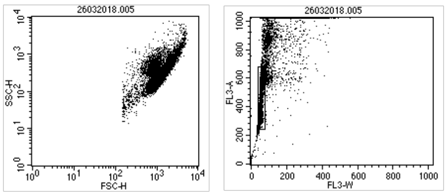


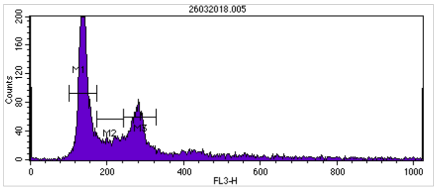


Histogram Statistics

File: 26032018.005
Sample ID:
Panel: Untitled Acquisition Tube List
Gate: No Gate
Total Events: 13495

Log Data Units: Linear Values
Tube: Untitled
Acquisition Date: 26-Mar-18
Gated Events: 13495
X Parameter: FL3-H (Linear)

| Marker | Left, Right | Events | % Gated | % Total | Mean | Geo Mean | CV | Median | Peak Ch |
|---|---|---|---|---|---|---|---|---|---|
| All | 0, 1023 | 13495 | 100.00 | 100.00 | 250.58 | 192.25 | 73.90 | 180.00 | 139 |
| M1 | 101, 173 | 6185 | 45.83 | 45.83 | 139.72 | 139.16 | 8.98 | 138.00 | 139 |
| M2 | 173, 243 | 1668 | 12.36 | 12.36 | 208.15 | 207.03 | 10.33 | 209.00 | 174 |
| M3 | 243, 328 | 3010 | 22.30 | 22.30 | 279.73 | 279.07 | 6.89 | 279.00 | 280 |

#### Chick control

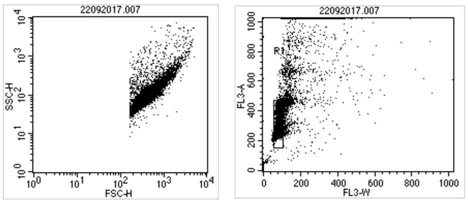


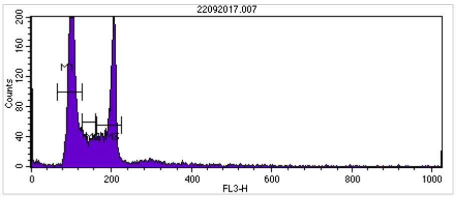


Histogram Statistics

File: 22092017.007
Sample ID:
Panel: Untitled Acquisition Tube List
Gate: No Gate
Total Events: 14204

Log Data Units: Linear Values
Tube: Untitled
Acquisition Date: 24-Oct-17
Gated Events: 14204
X Parameter: FL3-H (Linear)

| Marker | Left, Right | Events | % Gated | % Total | Mean | Geo Mean | CV | Median | Peak Ch |
|---|---|---|---|---|---|---|---|---|---|
| All | 0, 1023 | 14204 | 100.00 | 100.00 | 147.12 | 115.09 | 66.41 | 109.00 | 102 |
| M1 | 65, 127 | 7593 | 53.46 | 53.46 | 102.20 | 101.90 | 7.69 | 101.00 | 102 |
| M2 | 127, 160 | 1193 | 8.40 | 8.40 | 143.01 | 142.65 | 7.09 | 143.00 | 128 |
| M3 | 163, 225 | 3860 | 27.18 | 27.18 | 196.88 | 196.36 | 7.11 | 202.00 | 206 |

#### Quail polarising region cells grafted to chick

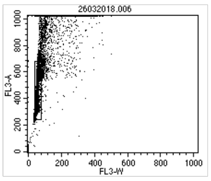


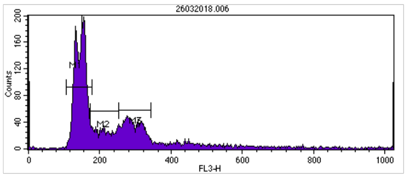


Histogram Statistics

File: 26032018.006
Sample ID:
Panel: Untitled Acquisition Tube List
Gate: No Gate
Total Events: 13102

Log Data Units: Linear Values
Tube: Untitled
Acquisition Date: 26-Mar-18
Gated Events: 13102
X Parameter: FL3-H (Linear)

| Marker | Left, Right | Events | % Gated | % Total | Mean | Geo Mean | CV | Median | Peak Ch |
|---|---|---|---|---|---|---|---|---|---|
| All | 0, 1023 | 13102 | 100.00 | 100.00 | 241.80 | 189.33 | 71.11 | 168.00 | 0 |
| M1 | 106, 178 | 6629 | 50.60 | 50.60 | 146.26 | 145.58 | 9.57 | 147.00 | 151 |
| M2 | 173, 253 | 1853 | 14.14 | 14.14 | 211.89 | 210.47 | 11.53 | 211.00 | 174 |
| M3 | 253, 344 | 2756 | 21.03 | 21.03 | 293.22 | 292.25 | 8.17 | 291.00 | 276 |

#### Chick polarising region grafted to a quail

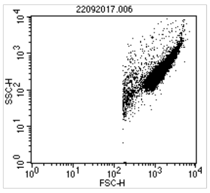


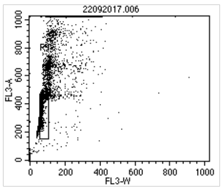


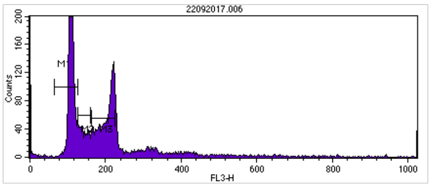


Histogram Statistics

File: 22092017.006
Sample ID:
Panel: Untitled Acquisition Tube List
Gate: No Gate
Total Events: 12772

Log Data Units: Linear Values
Tube: Untitled
Acquisition Date: 24-Oct-17
Gated Events: 12772
X Parameter: FL3-H (Linear)

| Marker | Left, Right | Events | % Gated | % Total | Mean | Geo Mean | CV | Median | Peak Ch |
|---|---|---|---|---|---|---|---|---|---|
| All | 0, 1023 | 12772 | 100.00 | 100.00 | 175.20 | 122.24 | 75.51 | 132.00 | 0 |
| M1 | 65, 127 | 5441 | 42.60 | 42.60 | 109.69 | 109.52 | 5.59 | 109.00 | 108 |
| M2 | 127, 160 | 1168 | 9.15 | 9.15 | 142.37 | 142.03 | 6.91 | 141.00 | 128 |
| M3 | 163, 225 | 3418 | 26.76 | 26.76 | 201.82 | 200.94 | 9.12 | 207.00 | 222 |

### 12hr polarising region grafts

#### Chick control

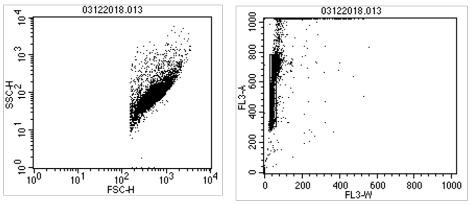


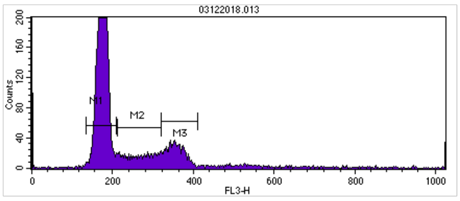


Histogram Statistics

File: 03122018.013
Sample ID:
Panel: Untitled Acquisition Tube List
Gate: No Gate
Total Events: 11730

Log Data Units: Linear Values
Tube: Untitled
Acquisition Date: 18-Mar-19
Gated Events: 11730
X Parameter: FL3-H (Linear)

| Marker | Left, Right | Events | % Gated | % Total | Mean | Geo Mean | CV | Median | Peak Ch |
|---|---|---|---|---|---|---|---|---|---|
| All | 0, 1023 | 11730 | 100.00 | 100.00 | 230.24 | 184.23 | 57.22 | 182.00 | 0 |
| M1 | 134, 209 | 7570 | 64.54 | 64.54 | 174.68 | 174.25 | 6.98 | 175.00 | 179 |
| M2 | 212, 320 | 1464 | 12.48 | 12.48 | 268.28 | 266.28 | 12.10 | 269.00 | 320 |
| M3 | 320, 411 | 1652 | 14.08 | 14.08 | 355.02 | 354.45 | 5.72 | 354.00 | 342 |

#### Quail control

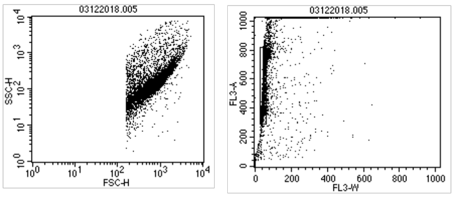


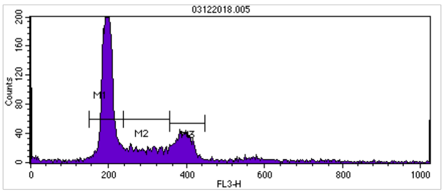


Histogram Statistics

File: 03122018.005
Sample ID:
Panel: Untitled Acquisition Tube List
Gate: No Gate
Total Events: 12885

Log Data Units: Linear Values
Tube: Untitled
Acquisition Date: 22-May-19
Gated Events: 12885
X Parameter: FL3-H (Linear)

| Marker | Left, Right | Events | % Gated | % Total | Mean | Geo Mean | CV | Median | Peak Ch |
|---|---|---|---|---|---|---|---|---|---|
| All | 0, 1023 | 12885 | 100.00 | 100.00 | 249.81 | 140.97 | 65.82 | 204.00 | 0 |
| M1 | 150, 238 | 6677 | 51.82 | 51.82 | 197.70 | 197.23 | 6.88 | 197.00 | 197 |
| M2 | 238, 357 | 1764 | 13.69 | 13.69 | 298.04 | 295.95 | 11.78 | 299.00 | 256 |
| M3 | 357, 447 | 1985 | 15.41 | 15.41 | 392.97 | 392.48 | 5.02 | 392.00 | 382 |

Quail polarising region grafted to chick host

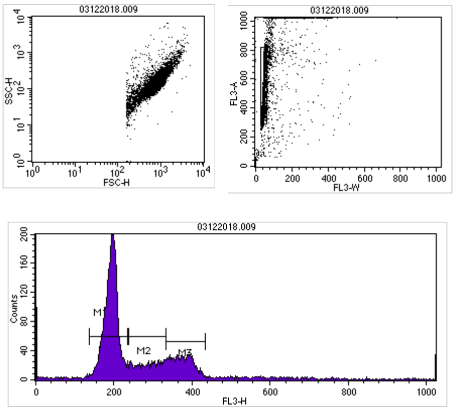


Histogram Statistics

File: 03122018.009
Sample ID:
Panel: Untitled Acquisition Tube List
Gate: No Gate
Total Events: 11697

Log Data Units: Linear Values
Tube: Untitled
Acquisition Date: 22-May-19
Gated Events: 11697
X Parameter: FL3-H (Linear)

| Marker | Left, Right | Events | % Gated | % Total | Mean | Geo Mean | CV | Median | Peak Ch |
|---|---|---|---|---|---|---|---|---|---|
| All | 0, 1023 | 11697 | 100.00 | 100.00 | 254.53 | 200.97 | 52.92 | 204.00 | 0 |
| M1 | 137, 235 | 6657 | 56.91 | 56.91 | 192.27 | 191.53 | 8.70 | 194.00 | 196 |
| M2 | 238, 333 | 1719 | 14.70 | 14.70 | 287.24 | 285.82 | 9.90 | 289.00 | 315 |
| M3 | 333, 434 | 2148 | 18.36 | 18.36 | 374.52 | 373.71 | 6.60 | 374.00 | 343 |

Chick polarising region to quail host

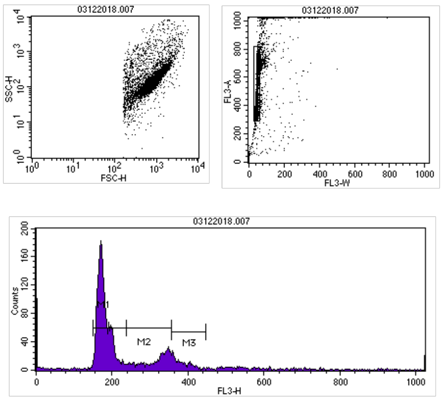


Histogram Statistics

File: 03122018.007
Sample ID:
Panel: Untitled Acquisition Tube List
Gate: No Gate
Total Events: 8250

Log Data Units: Linear Values
Tube: Untitled
Acquisition Date: 22-May-19
Gated Events: 8250
X Parameter: FL3-H (Linear)

| Marker | Left, Right | Events | % Gated | % Total | Mean | Geo Mean | CV | Median | Peak Ch |
|---|---|---|---|---|---|---|---|---|---|
| All | 0, 1023 | 8250 | 100.00 | 100.00 | 211.85 | 110.69 | 69.44 | 179.00 | 0 |
| M1 | 150, 238 | 4849 | 58.78 | 58.78 | 178.87 | 178.15 | 9.22 | 175.00 | 171 |
| M2 | 238, 357 | 1254 | 15.20 | 15.20 | 314.17 | 312.03 | 11.31 | 327.00 | 348 |
| M3 | 357, 447 | 577 | 6.99 | 6.99 | 382.62 | 382.04 | 5.54 | 378.00 | 358 |

| 0hr grafts | | | | | |
|---|---|---|---|---|---|
| | | Quail control | Chick control | Quail graft to chick | Chick graft to quail |
| **Number of cells:** | G1 | 6185 | 7593 | 6629 | 5441 |
| | S/G2/M | 4678 | 5053 | 4609 | 4586 |

*p*-values:

Quail vs Quail grafted to Chick = 0.02

Chick vs Quail grafted to Chick = 0.09

Chick vs Chick grafted to Quail = 0.00001

Quail vs Chick grafted to Quail = 0.00012

| 12hr grafts | | | | | |
|---|---|---|---|---|---|
| | | Quail control | Chick control | Quail graft to chick | Chick graft to quail |
| **Number of cells:** | G1 | 6677 | 7570 | 6657 | 4849 |
| | S/G2/M | 3749 | 3116 | 3867 | 1831 |

*p*-values:

Quail vs Quail grafted to Chick = 0.24

Chick vs Quail grafted to Chick = 0.00001

Chick vs Chick grafted to Quail = 0.013

Quail vs Chick grafted to Quail = 0.00001

1. Scherz PJ, Harfe BD, McMahon AP, Tabin CJ. The limb bud Shh-Fgf feedback loop is terminated by expansion of former ZPA cells. Science. 305. United States2004. p. 396-9.
